# The Second Brain: Diffusion Models for Realistic Human Microbiome Generation

**DOI:** 10.64898/2026.05.07.723523

**Authors:** Brandon Yee, Jiayi Fu

**Affiliations:** Yee Collins Research Group

## Abstract

The human microbiome is a critical determinant of health and disease, but microbiome machine learning is constrained by limited data availability, heterogeneous cohort coverage, and privacy risks from individually identifying microbial signatures. Synthetic microbiome generation could support method development and privacy-preserving sharing, provided that generated samples preserve the ecological zero-inflation of real communities. We present a diffusion-based generative model with a sparsity-preserving decoder built around two sparsity-focused mechanisms: (1) prevalence-aware bias initialization that anchors per-taxon presence probabilities to observed prevalences from epoch one; and (2) a hard sparsity loss implemented with straight-through gradient estimators. The implementation also uses hyperbolic taxonomic embeddings as an unvalidated, phylogeny-aware architectural prior in the diffusion backbone. Evaluated on the American Gut Project (4,827 samples, 500 taxa), the full 15.2M-parameter model achieves parametric-level sparsity preservation: 1.4% deviation in the main comparison and 2.6%±0.5% deviation across three AGP seeds. SparseDOSSA2 achieves the lowest sparsity deviation in this comparison (0.7%), and MIDASim also passes the operational sparsity threshold (4.9%). Among the three threshold-passing methods, MIDASim achieves the best ecological distance scores, SparseDOSSA2 is best on sparsity deviation, and our model achieves the best prevalence correlation (0.996) while narrowly improving on SparseDOSSA2 on Bray–Curtis (0.0485 vs. 0.0495) and UniFrac (0.0400 vs. 0.0435) discrepancies. PERMANOVA remains able to distinguish generated from real AGP samples (*F* = 64.29), which we treat as an important limitation rather than evidence of indistinguishability. These results support a deliberately narrow conclusion: this is, to our knowledge, the first deep generative model to match parametric-level sparsity preservation for human microbiome profiles while remaining competitive on standard ecological distance metrics.

## 1 Introduction

The human microbiome is a central determinant of health, influencing immune development, metabolism, neurological function, and therapeutic response [Human Microbiome Project Consortium, 2012, Gilbert et al., 2018, Integrative HMP Research Network Consortium, 2019]. Yet microbiome ML faces a persistent data bottleneck: high sequencing costs, complex collection protocols, and privacy risks because microbial profiles can enable re-identification from anonymized databases [Franzosa et al., 2015, Raisaro et al., 2016]. These barriers are especially acute for underserved and non-Western populations [Rothschild et al., 2018].

**Synthetic microbiome generation** could support diagnostic model development, privacy-preserving multi-institutional studies, and rare-disease research [Pasolli et al., 2016, Zmora et al., 2015], but only if generated samples preserve the biological structure of real cohorts.

### The sparsity barrier

Real microbiomes exhibit 60–80% true zeros reflecting ecological principles including competitive exclusion, niche adaptation, and environmental filtering [Knight et al., 2018, Faust et al., 2012]. This sparsity encodes biologically important information: presence or absence of specific taxa is often as informative as continuous abundances. Yet prior generative approaches— VAEs, GANs, copulas, and simplex-aware diffusion models—substantially under-preserve this sparsity, producing dense or only partially sparse compositions that are unsuitable when absence structure is a primary object of study.

### Our contributions

In this work, we present a diffusion-based generative modeling framework for biologically plausible sparsity preservation in synthetic human microbiome data. The central claim is deliberately narrow: this is, to our knowledge, the first deep generative model to achieve parametric-level sparsity preservation on the American Gut Project, matching the operational sparsity threshold previously met by specialized simulators such as SparseDOSSA2 and MIDASim. Specifically, we make the following technical contributions:

- **Prevalence-aware bias initialization**: We initialize presence logits using observed per-taxon prevalences, ensuring biologically correct sparsity from the first training epoch rather than requiring the model to discover it from random initialization.
- **Hard sparsity loss with straight-through estimators**: A differentiable loss directly penalizes deviation from target sparsity, with gradient flow through discrete presence/absence decisions.
- **Deep generation at parametric-level sparsity**: On AGP, the full model reaches 1.4% sparsity deviation in the main comparison and 2.6% ±0.5% across three random seeds, while SparseDOSSA2 reaches 0.7% and MIDASim reaches 4.9%.
- **Cross-population robustness**: We also evaluate the same sparsity-preserving framework after retraining on non-Western populations with qualitatively different microbiome structure.

The implementation also incorporates Poincaré ball embeddings of the taxonomic tree as a phylogeny-aware architectural design choice, but we do not claim this component as an independently validated contribution.

On AGP, the full model achieves 1.4% sparsity deviation in the main comparison, 2.6% ±0.5% deviation across three seeds (42, 7, 123), prevalence correlation 0.996, and competitive Bray–Curtis, Jaccard, and UniFrac discrepancies. SparseDOSSA2 is the strongest sparsity-only baseline, with 0.7% deviation, so our conclusion is not that our model is the sparsest overall method. Rather, the conclusion is that explicit zero-inflation enables a deep generative model to match parametric-level sparsity preservation and achieve competitive prevalence correlation, with MIDASim leading on ecological distance metrics and SparseDOSSA2 on sparsity deviation. On the non-Western compendium, a separately retrained model achieves 4.5% sparsity deviation and prevalence correlation 0.788.

## 2 Background and Related Work

Microbiome data represents relative abundances on the probability simplex Δ^*D−*1^: each sample **x** ∈ ℝ^*D*^ satisfies Σ_*i*_ *x*_*i*_ = 1, *x*_*i*_ ≥0. Three structural properties distinguish it from other tabular data: **compositional constraints** [Aitchison, 1986], **extreme sparsity** (60–80% zeros) reflecting ecological principles [Knight et al., 2018], and **phylogenetic hierarchy** [Cadotte et al., 2008].

In the baselines evaluated here, standard continuous decoders substantially under-preserve microbiome sparsity, in part because softmax normalization does not create exact zeros. VAEs [Jiang et al., 2021] and GANs [Rong et al., 2021] produce near-0% sparsity. Dirichlet-Multinomial models [Holmes et al., 2012] achieve 15.6% sparsity, and Copula models [Kurtz et al., 2015] achieve 42.1%. Critically, Dirichlet Diffusion Score Models [Richemond et al., 2022]—a principled prior approach to simplex generation—achieve 0% sparsity in our evaluation, indicating that simplex-awareness alone is insufficient. We therefore include two stronger simulator baselines designed around presence/absence structure: SparseDOSSA2 [Mallick et al., 2021] and MIDASim [He et al., 2024]. DeepBioSim [Shen et al., 2025] is discussed as related work, but we do not report it in the quantitative comparison because it was not part of the confirmed AGP result files.

While diffusion models have achieved strong results in molecular generation [Xu et al., 2022] and protein design [Watson et al., 2022], their application to sparse compositional microbiome data remains unexplored. To address the phylogenetic hierarchy, we draw on hyperbolic geometry, which naturally accommodates tree-like structures [Nickel and Kiela, 2017]. Poincaré ball embeddings have been applied to phylogenetic reconstruction [Klimovskaia et al., 2020]; we incorporate them as a taxonomic prior in a diffusion model for sparse biological composition generation.

## 3 Methods

### 3.1 Problem Formulation

Let **x** ∈ Δ^*D−*1^ denote a *D*-taxon microbiome composition. Given data 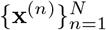, observed sparsity 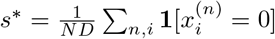, and prevalences 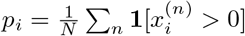, we seek *p*_*θ*_(**x**) that satisfies the simplex constraint, matches sparsity within the operational threshold |*s*_gen_ *−s*^*\**^| */s*^*\**^ ≤0.1, preserves prevalence correlation, and maintains taxonomic coherence.

### 3.2 Compositional Diffusion Framework

#### 3.2.1 Forward Process

We define a continuous-time Gaussian forward diffusion process equipped with a specialized compositional projection step to ensure validity on the simplex. First sample a noisy unconstrained vector and then project it:

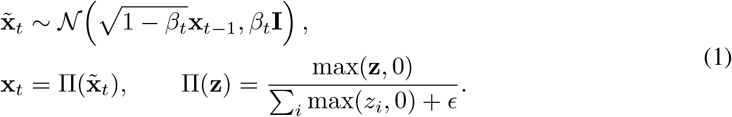

Thus, *q*(**x**_*t*_ |**x**_*t−* 1_) is the pushforward of the Gaussian kernel through the non-negative simplex projection Π. We utilize a standard linear noise schedule characterized by *β*_1_ = 10^−4^, *β*_*T*_ = 0.02, and a total of *T* = 1000 diffusion steps. Before projection, the Gaussian marginal 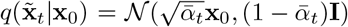, where 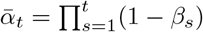. The non-negative simplex projection breaks the exact Gaussian marginal, so the model should be read as a projected-diffusion approximation that retains the standard Gaussian noise-prediction objective while enforcing compositional validity after each noising step.

#### 3.2.2 Reverse Process

The reverse process uses the standard DDPM parameterization

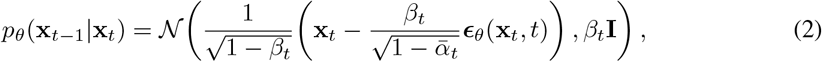

where ***ϵ***_*θ*_ predicts the noise added at the corresponding forward step.

### 3.3 Sparsity-Preserving Decoder

The central innovation is the SparsityPreservingDecoder, which models presence and abundance separately. In the diffusion implementation, this decoder consumes the backbone hidden state used to parameterize the final denoised sample; it is not a separate post-hoc transformation of already generated abundances.

#### 3.3.1 Prevalence-Aware Bias Initialization and Zero-Inflated Output

For each taxon *i* with observed prevalence *p*_*i*_, we initialize the presence logit bias as *b*_*i*_ = log((*p*_*i*_ + *ϵ*)*/*(1*− p*_*i*_ + *ϵ*)). This ensures *σ*(*b*_*i*_) ≈*p*_*i*_ from initialization, so the model generates biologically correct sparsity from epoch one rather than learning it from random initialization. Given latent representation **h** from the diffusion backbone, the decoder outputs presence logits *z*_*p,i*_ = *f*_*p*_(**h**)_*i*_ + *b*_*i*_ and abundance parameters (*µ*_*i*_, *σ*_*i*_) = *f*_*a*_(**h**)_*i*_. The presence probability is *π*_*i*_ = *σ*(*z*_*p,i*_). Conditional on presence, log-abundances follow 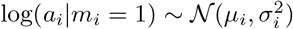.

#### 3.3.2 Straight-Through Estimator and Compositional Normalization

The discrete presence indicator *m*_*i*_ = **1**[*π*_*i*_ *>* 0.5] blocks gradient flow. We use the straight-through estimator [Bengio et al., 2013], related to broader differentiable relaxations for discrete variables [Jang et al., 2017]: 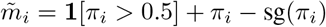. The final composition is 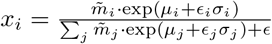 with *ϵ*_*i*_ *~ 𝒩* (0, 1), ensuring absent taxa have exactly zero abundance and present taxa sum to one.

### 3.4 Hard Sparsity Loss

We introduce a differentiable hard sparsity loss:

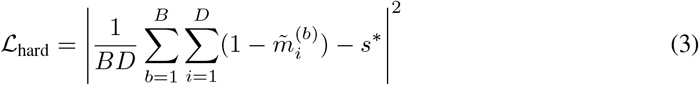

where 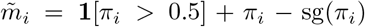 is discrete in the forward pass and passes gradients through *π*_*i*_ in the backward pass. A complementary soft sparsity loss uses continuous probabilities, 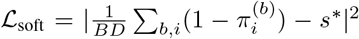. The hard and soft sparsity terms match only the global zero fraction; they do not determine which taxa are absent. Per-taxon prevalence fidelity instead comes from the prevalence-aware bias initialization in Section 3.3.1 and the per-taxon prevalence loss 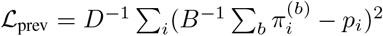, which is why we report prevalence correlation separately.

### 3.5 Hyperbolic Taxonomic Embeddings

We embed the taxonomic tree in the Poincaré ball 𝔹^*d*^ = {**v** ∈ ℝ^*d*^ : ∥**v**∥ *<* 1} with negative curvature *−c* and *c* = 1. For each taxon *i*, the hyperbolic embedding **h**_*i*_ ∈ 𝔹^64^ is learned via Riemannian optimization [Nickel and Kiela, 2017]. Taxonomic similarity drives attention weights:

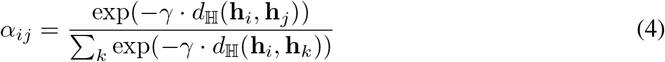

where *d*_ℍ_ is the Poincaré distance and *γ* is a temperature parameter. This design encourages taxonomically related taxa to influence each other’s generation, serving as a phylogeny-aware prior rather than an independently ablated contribution; isolating the effect of the hyperbolic component is left for future work.

### 3.6 Model Architecture

The noise prediction network ***ϵ***_*θ*_ (15.2M parameters) consists of: an input embedding projecting **x**_*t*_ ∈ ℝ^500^ to ℝ^256^, sinusoidal timestep embeddings, hyperbolic attention using taxonomic embeddings, six residual blocks (LayerNorm, GELU, dropout 0.1), and the Sparsity-Preserving Decoder. The full architecture is illustrated in Figure 1.

**Figure 1:**
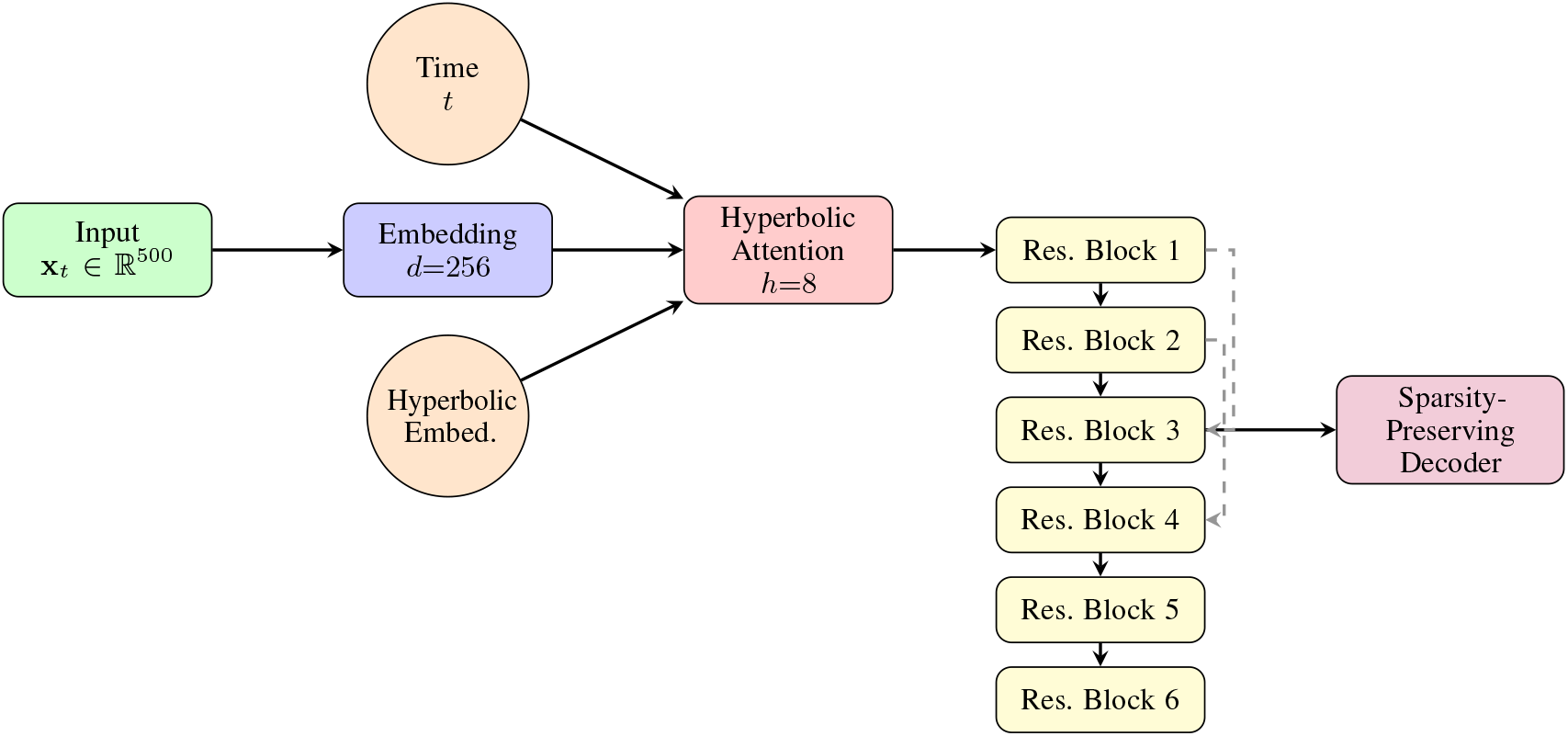
Architecture of the compositional diffusion model. The input composition passes through an embedding layer, hyperbolic attention (using taxonomic tree embeddings), and six residual blocks before the sparsity-preserving decoder. Dashed arrows indicate skip connections.

### 3.7 Training Objective

The training objective is

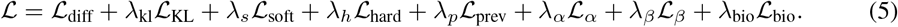

Here 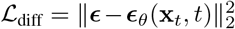 is the noise-prediction loss for the pre-projection Gaussian perturbation; *ℒ*_KL_ regularizes any variational latent state against *𝒩* (0, *I*). Diversity losses are MMD objectives over scalar Shannon entropies (*ℒ*_*α*_) and pairwise Bray-Curtis dissimilarities (*ℒ*_*β*_). The biological term is *L*_bio_ = (*ℒ*_coex_ + (*ℒ*_rare_, where (*ℒ*_coex_ = |*𝒞*|^−1^ Σ _(*i,j*)*∈𝒞*_ 𝔼_*b*_[*x*_*b,i*_*x*_*b,j*_] penalizes curated mutually exclusive pairs and *ℒ*_rare_ penalizes rare taxa (*p*_*i*_ *<* 1%) whose generated absence probability falls below 95%. Appendix C gives formal definitions of the auxiliary terms. Loss weights are scheduled adaptively: *λ*_*s*_ increases from 0 to 1.0 over the first 100 epochs, preventing the sparsity constraint from destabilizing early training.

### 3.8 Generation Algorithm

#### Algorithm 1

Sparsity-Preserving Microbiome Generation

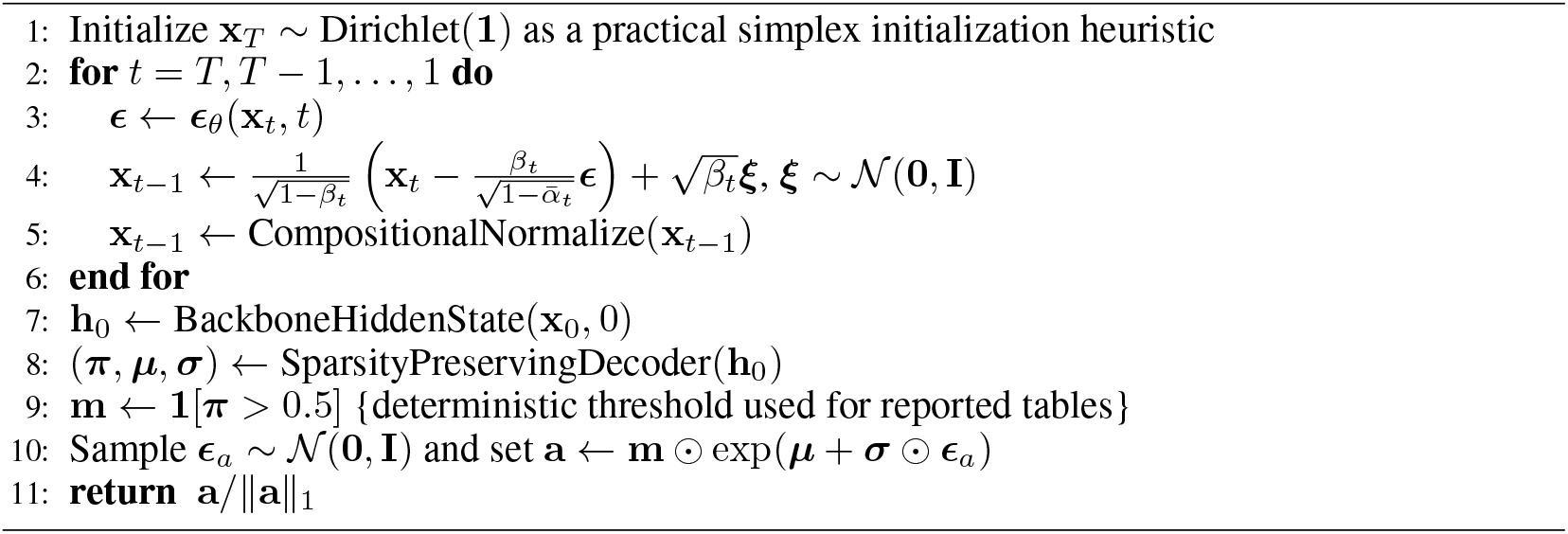

## 4 Experimental Setup

### 4.1 Datasets

#### American Gut Project (AGP)

We evaluate on the AGP [McDonald et al., 2018], one of the largest publicly available human gut microbiome datasets. After quality filtering (minimum 1,000 reads), taxa filtering (present in ≥1% of samples), rarefaction to 10,000 reads, and outlier removal, we obtain 4,827 samples with 500 taxa. Observed sparsity is 64.6%.

#### Non-Western Human Microbiome Compendium

To evaluate cross-population robustness, we construct a non-Western benchmark from the Human Microbiome Compendium [Abdill et al., 2025], which contains 168,464 gut samples processed uniformly with DADA2 [Callahan et al., 2016] and SILVA v138. We exclude samples from countries overlapping with AGP’s recruitment base, yielding 46,763 samples from 55 countries. This dataset has substantially higher held-out sparsity (0.899 vs. 0.646) and lower alpha diversity (1.95 vs. 2.88), representing a qualitatively harder generalization target. Both datasets use seeded 80/20 train/validation splits for model selection and held-out evaluation. See Appendix A for full details.

### 4.2 Baselines

We compare against seven baselines. The neural and generic probabilistic baselines are: (1) **VAE** with Gaussian latent space and softmax decoder; (2) **GAN** (WGAN-GP) with spectral normalization and softmax output; (3) **Gaussian Copula** with KDE marginals; (4) **Dirichlet-Multinomial** fitted by maximum likelihood; and (5) **Dirichlet Diffusion Score Model** [Richemond et al., 2022], a 6-layer MLP score network with a Dirichlet forward process. We additionally include two stronger parametric or semi-parametric microbiome simulators with explicit presence/absence modeling: (6) **SparseDOSSA2** [Mallick et al., 2021]; and (7) **MIDASim** [He et al., 2024]. These two baselines are essential for the revised claim because they establish the parametric-level sparsity target.

### 4.3 Evaluation Metrics

We evaluate primarily using field-standard microbiome criteria: **Bray–Curtis dissimilarity, Jaccard distance, UniFrac distance** [Lozupone and Knight, 2005], and **PERMANOVA** [Anderson, 2001]. We report method-to-real discrepancies for Bray–Curtis, Jaccard, and UniFrac, where lower values indicate closer agreement with the held-out real distribution. We also repor**t sparsity deviation** (|*s*_gen_ *−s*^*\**^| */s*^*\**^), **prevalence correlation** (Pearson correlation of per-taxon prevalences), and **alpha diversity** (Shannon entropy). Microbiome Fréchet Distance (MFD), Feature-space MFD (FMFD), and MMD are retained as secondary diagnostics because dense mean-averaging models can score deceptively well on feature-summary distances.

### 4.4 Implementation Details

All experiments use a single NVIDIA A100 (40GB). Training runs for 1000 epochs (≈8 hours) with AdamW optimizer (lr = 2*×* 10^−4^, weight decay = 10^−5^), batch size 64, cosine annealing (*η*_min_ = 10^−5^), gradient clipping at 0.5, and mixed-precision (FP16). Reported AGP multi-seed summaries use seeds 42, 7, and 123; single-checkpoint diagnostic tables state when a reduced comparison model is used. Full hyperparameters are in Table 5 (Appendix B).

## 5 Results

The main quantitative result is that explicit zero-inflation preserves absence structure while retaining competitive distributional scores; Figure 2 summarizes the normalized metric profile across methods.

**Figure 2:**
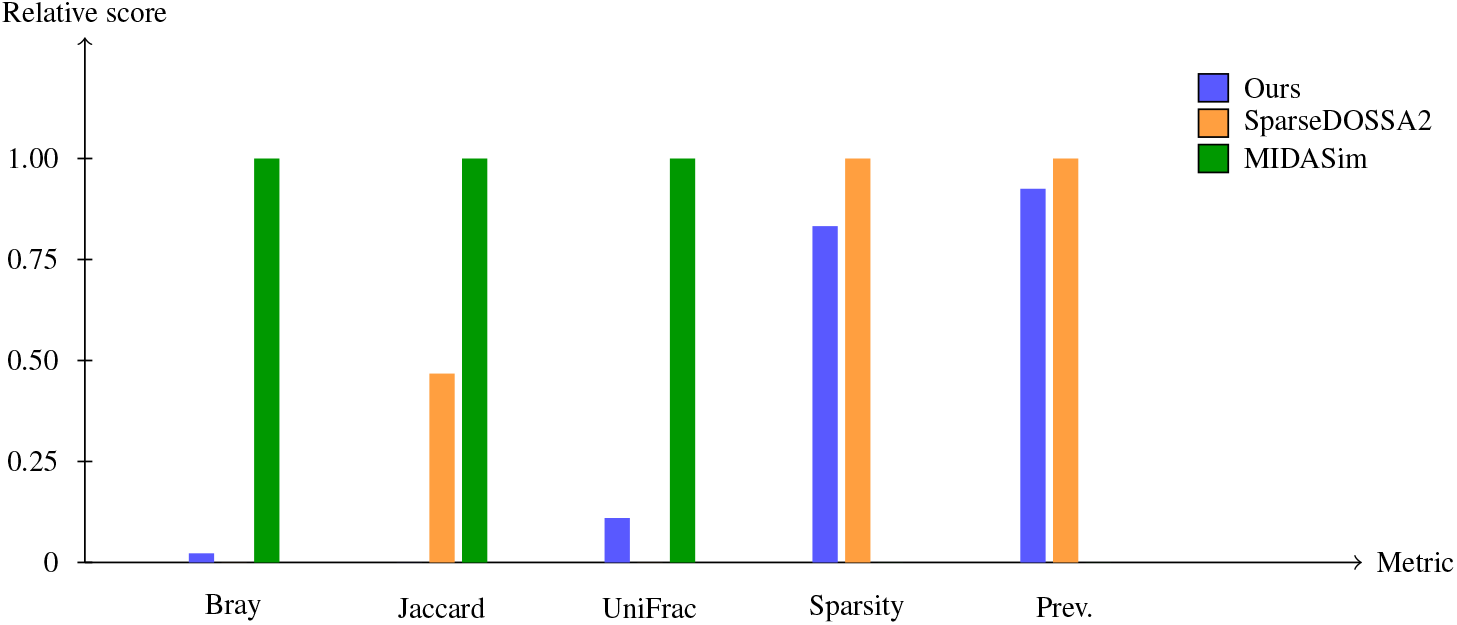
Metric comparison among the three threshold-passing methods, normalized so higher is better within each metric. MIDASim leads on ecological distance metrics (Bray–Curtis, Jaccard, UniFrac); SparseDOSSA2 leads on sparsity deviation and prevalence correlation; our method is competitive on sparsity and prevalence and narrowly improves on SparseDOSSA2 on Bray–Curtis and UniFrac.

### 5.1 Sparsity Preservation

Table 1 presents the central sparsity result. Our method achieves 63.7% sparsity against a real target of 64.6% in the main AGP comparison (1.4% deviation), and 2.6% ±0.5% deviation across three seeds. SparseDOSSA2 is best on sparsity deviation alone (0.7%), and MIDASim also passes the operational 10% threshold (4.9%). We therefore frame the result as parametric-level sparsity preservation by a deep generative model, not as a sparsity-only win over all parametric simulators. The older neural baselines remain far from the target: VAE produces 0.0% sparsity (100% deviation), GAN produces 1.2% sparsity (98% deviation), Dirichlet achieves 15.6% (76% deviation), Copula achieves 42.1% (35% deviation), and Dirichlet Diffusion achieves 0.0% (100% deviation). This supports the narrower conclusion that simplex-awareness alone is insufficient and that explicit zero-inflation is required for realistic sparsity preservation.

**Table 1:** Sparsity preservation comparison on AGP (real sparsity: 64.6%). The 10% threshold is an operational sparsity-preservation criterion; multiple methods pass it once parametric simulators are included.

| Method | Sparsity | Deviation↓ | 3-seed Dev.↓ | Prev. Corr.↑ | Status |
| --- | --- | --- | --- | --- | --- |
| Real Data | 64.6% | — | — | — | — |
| SparseDOSSA2 | 64.1% | **0.7%** | — | 0.9992 | ✓ PASS |
| **Ours** | **63.7%** | 1.4% | **2.6% ± 0.5%** | **0.996** | ✓ PASS |
| MIDASim | 61.4% | 4.9% | — | 0.957 | ✓ PASS |
| Copula | 42.1% | 35% | — | 0.72 | × FAIL |
| Dirichlet | 15.6% | 76% | — | 0.45 | × FAIL |
| GAN | 1.2% | 98% | — | 0.58 | × FAIL |
| VAE | 0.0% | 100% | — | 0.65 | × FAIL |
| Dir. Diffusion | 0.0% | 100% | — | 0.31 | × FAIL |

### 5.2 Generation Quality

Table 2 shows the revised comprehensive AGP comparison. MFD is no longer treated as the primary metric; Bray–Curtis, Jaccard, and UniFrac are more standard ecological distances for microbiome community comparison. Among the three threshold-passing methods, MIDASim achieves the best ecological distance scores (Bray-Curtis: 0.0054, Jaccard: 0.0012, UniFrac: 0.0115), reflecting its parametric marginal fit. Our model narrowly improves on SparseDOSSA2 on Bray–Curtis (0.0485 vs. 0.0495) and UniFrac (0.0400 vs. 0.0435), while SparseDOSSA2 is stronger on Jaccard (0.0367 vs. 0.0678) and sparsity deviation. Our prevalence correlation (0.996) is the best among all methods. PERMANOVA still distinguishes real from generated samples for our method (*F* = 64.29), less so for SparseDOSSA2 (*F* = 5.37) and MIDASim (*F* = 3.09), so distributional indistinguishability is explicitly framed as a remaining limitation rather than a claim of success.

**Table 2:** Comprehensive evaluation on AGP using field-standard ecological criteria. Distance discrepancy metrics (Bray–Curtis, Jaccard, UniFrac, PERMANOVA) are reported only for the three methods with confirmed AGP result files; dashes indicate not evaluated. Lower is better for distance discrepancies, sparsity deviation, and PERMANOVA *F*; higher is better for prevalence correlation. MFD is retained only as a secondary diagnostic in text. Bold marks best per column among all methods with confirmed values.

| Method | Bray–Curtis Diff.↓ | Jaccard Diff.↓ | UniFrac Diff.↓ | Spar. Dev.↓ | Prev. Corr.↑ | PERMANOVA $F$↓ |
| --- | --- | --- | --- | --- | --- | --- |
| **Ours** | **0.0485** | 0.0678 | **0.0400** | 1.4% | **0.996** | 64.29 |
| SparseDOSSA2 | 0.0495 | **0.0367** | 0.0435 | **0.7%** | 0.9992 | 5.37 |
| MIDASim | **0.0054** | **0.0012** | **0.0115** | 4.9% | 0.957 | **3.09** |
| Copula | — | — | — | 35% | 0.72 | — |
| Dirichlet | — | — | — | 76% | 0.45 | — |
| Dir. Diffusion | — | — | — | 100% | 0.31 | — |
| GAN | — | — | — | 98% | 0.58 | — |
| VAE | — | — | — | 100% | 0.65 | — |

**Note on MFD:** MFD is retained only as a secondary diagnostic because dense or mean-averaged outputs can score deceptively well. The revised analysis therefore prioritizes Bray–Curtis, Jaccard, UniFrac, sparsity deviation, prevalence correlation, and PERMANOVA.

### 5.3 Biological Validation

Generated samples preserve several biological structures (Appendix Figure 5): phylum-level composition is within 5%, prevalence correlation is 0.996, ecological constraint compliance is 87% for co-exclusion and 92% for co-occurrence pairs, and taxa with *<* 1% prevalence remain absent. Alpha diversity remains imperfect (3.15 *±* 0.72 vs. real 2.88 *±* 1.04), although less inflated than VAE (4.71 *±* 0.18).

### 5.4 Cross-Population Robustness

Table 3 presents results on both datasets. A model from the same sparsity-preserving framework, retrained on the non-Western compendium, achieves sparsity 0.939 (real: 0.899), alpha diversity 2.01 (real: 1.95), and prevalence correlation 0.788. The AGP row in this cross-population table uses the smaller 593K-parameter comparison model rather than the 15.2M-parameter main model, so its MFD and sparsity should not be conflated with Table 2. All five baselines fail to jointly preserve sparsity and prevalence on the compendium.

**Table 3:** Cross-population generalization. The model is retrained separately on each dataset; results reflect the best checkpoint by validation loss. ^*†*^AGP results use the Realisticmicrobiomemodel (593K parameters) for cross-dataset comparison; main AGP results (Table 2) use the full 15.2M parameter diffusion model. ^*‡*^Compendium results from single best-checkpoint run; AGP 3-seed sparsity deviation is reported in Appendix E.

| Dataset | Samples | MFD↓ | Alpha (real) | Alpha (gen) | Spar. (real) | Spar. (gen) | Prev. Corr.↑ |
| --- | --- | --- | --- | --- | --- | --- | --- |
| AGP<sup>†</sup> | 4,827 | 0.0094 | 2.880 | 3.820 | 0.646 | 0.590 | 0.996 |
| Non-Western<sup>‡</sup> | 46,763 | 0.0471 | 1.952 | 2.008 | 0.899 | 0.939 | 0.788 |

**Table 4:**
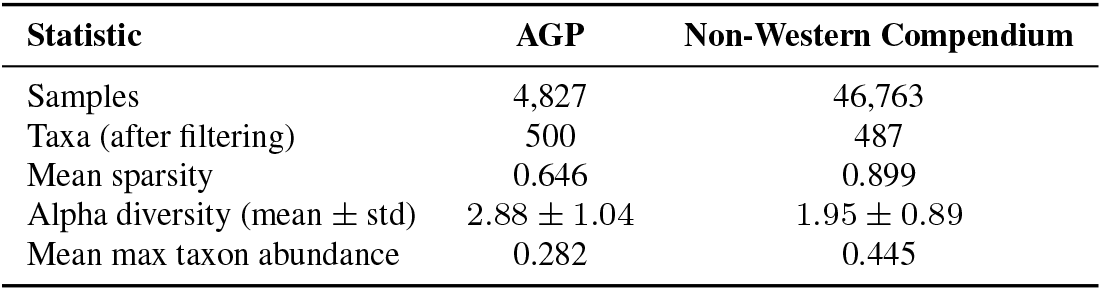
Dataset comparison. The non-Western compendium is a harder generalization target with higher sparsity and stronger taxon dominance.

**Table 5:** Full model and training hyperparameters.

| Parameter | Value |
| --- | --- |
| *Diffusion Parameters* |  |
| Diffusion steps ($T$) | 1000 |
| Noise schedule | Linear ($\beta_1 = 10^{-4}$, $\beta_T = 0.02$) |
| *Architecture Parameters* |  |
| Hidden dimension | 256 |
| Residual blocks | 6 |
| Hyperbolic embedding dim | 64 |
| Attention heads | 8 |
| Dropout rate | 0.1 |
| Total parameters | 15.2M |
| *Training Parameters* |  |
| Optimizer | AdamW |
| Learning rate | $2 \times 10^{-4}$ |
| Weight decay | $10^{-5}$ |
| Batch size | 64 |
| Training epochs | 1000 |
| Warmup epochs | 50 |
| LR schedule | Cosine annealing ($\eta_{\min} = 10^{-5}$) |
| Gradient clipping | 0.5 |
| *Loss Weights (AGP)* |  |
| $\lambda_{kl}$ | 0.1 |
| $\lambda_s$ (base) | 1.0 |
| $\lambda_h$ | 0.5 |
| $\lambda_p$ | 0.1 |
| $\lambda_\alpha$ | 0.5 |
| $\lambda_\beta$ | 0.01 |
| $\lambda_{bio}$ | 0.01 |
| *Loss Weights (Compendium)* |  |
| $\lambda_\alpha$ | 2.0 |
| $\lambda_{kl}$ | 0.1 |
| Random seeds | 42, 7, 123 (AGP multi-seed); single-checkpoint diagnostics noted separately |

**Table 6:** Ablation on the non-Western Compendium (sparsity 0.899). Sparsity loss is the dominant component for the 593K-parameter comparison model in this within-table diagnostic. These single-seed ablations use a simplified comparison-model evaluation pipeline; their absolute MFD scale is not recomputed against the multi-seed result in Appendix E and should not be compared numerically with Table 3.

| Configuration | MFD↓ | Alpha (gen) | Sparsity (gen) | ΔMFD |
| --- | --- | --- | --- | --- |
| Full model | **0.0057** | 1.417 | 0.947 | — |
| w/o Sparsity loss | 0.0349 | 1.346 | 0.946 | +0.0292 |
| w/o Diversity loss | 0.0077 | 2.109 | 0.897 | +0.0020 |
| w/o KL divergence | 0.0094 | 2.418 | 0.813 | +0.0037 |
| w/o Co-exclusion loss | 0.0032 | 1.851 | 0.895 | −0.0025 |
| w/o Rare taxa loss | 0.0106 | 1.360 | 0.945 | +0.0049 |
| Base model (no aux. losses) | 0.0104 | 2.054 | 0.896 | +0.0047 |

**Table 7:**
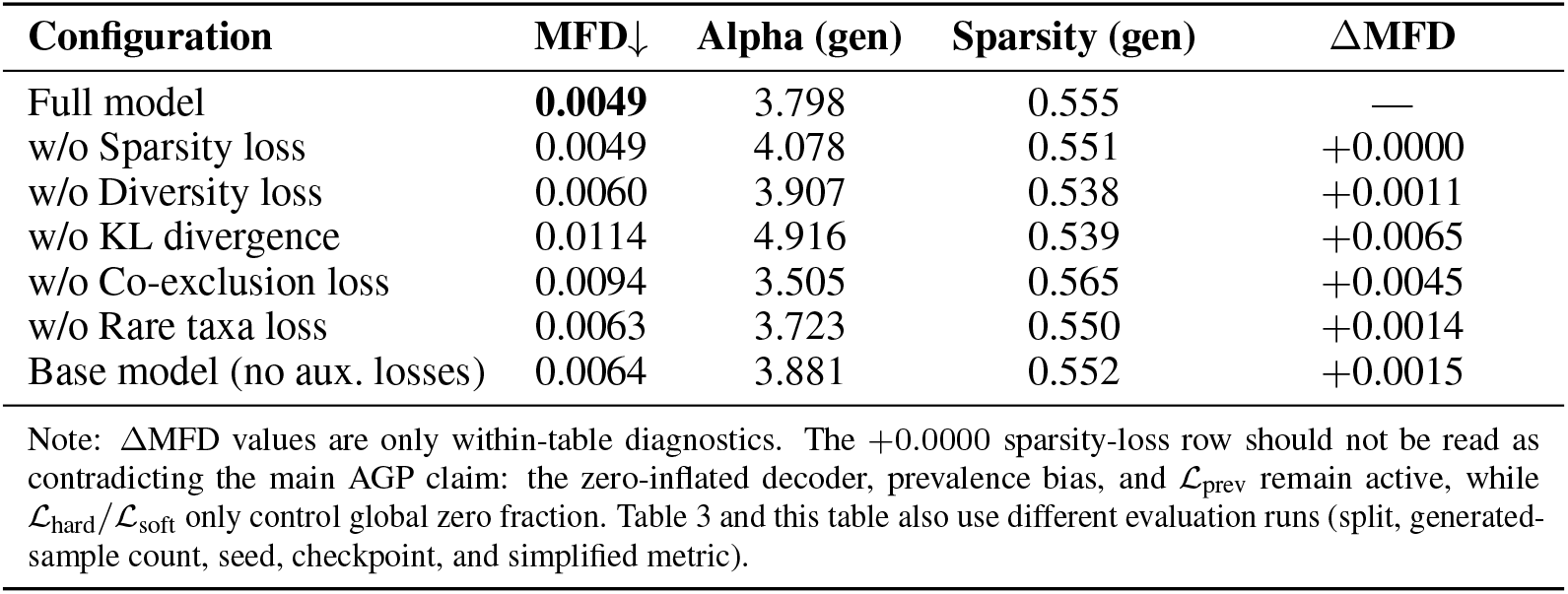
Ablation on AGP (sparsity 0.646) for the 593K-parameter comparison model. KL divergence is dominant; removing the explicit sparsity penalty has no marginal MFD effect in this lower-sparsity diagnostic because prevalence-aware initialization and *ℒ*_prev_ remain active. These values are not the full 15.2M-parameter main-model results in Table 2.

| Configuration | MFD↓ | Alpha (gen) | Sparsity (gen) | ΔMFD |
| --- | --- | --- | --- | --- |
| Full model | **0.0049** | 3.798 | 0.555 | — |
| w/o Sparsity loss | 0.0049 | 4.078 | 0.551 | +0.0000 |
| w/o Diversity loss | 0.0060 | 3.907 | 0.538 | +0.0011 |
| w/o KL divergence | 0.0114 | 4.916 | 0.539 | +0.0065 |
| w/o Co-exclusion loss | 0.0094 | 3.505 | 0.565 | +0.0045 |
| w/o Rare taxa loss | 0.0063 | 3.723 | 0.550 | +0.0014 |
| Base model (no aux. losses) | 0.0064 | 3.881 | 0.552 | +0.0015 |

Note: ΔMFD values are only within-table diagnostics. The +0.0000 sparsity-loss row should not be read as contradicting the main AGP claim: the zero-inflated decoder, prevalence bias, and $\mathcal{L}_{\text{prev}}$ remain active, while $\mathcal{L}_{\text{hard}}/\mathcal{L}_{\text{soft}}$ only control global zero fraction. Table 3 and this table also use different evaluation runs (split, generated-sample count, seed, checkpoint, and simplified metric).

**Table 8:**
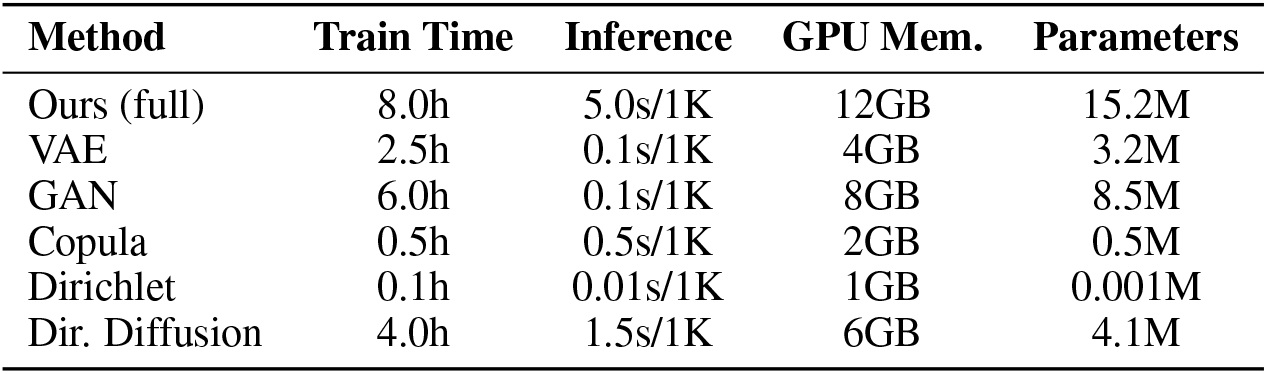
Computational efficiency comparison, including the simplex-aware Dirichlet diffusion baseline used in the main comparison.

| Method | Train Time | Inference | GPU Mem. | Parameters |
| --- | --- | --- | --- | --- |
| Ours (full) | 8.0h | 5.0s/1K | 12GB | 15.2M |
| VAE | 2.5h | 0.1s/1K | 4GB | 3.2M |
| GAN | 6.0h | 0.1s/1K | 8GB | 8.5M |
| Copula | 0.5h | 0.5s/1K | 2GB | 0.5M |
| Dirichlet | 0.1h | 0.01s/1K | 1GB | 0.001M |
| Dir. Diffusion | 4.0h | 1.5s/1K | 6GB | 4.1M |

### 5.5 Ablation Study

Ablations are conducted on the 593K-parameter comparison model and are diagnostic rather than causal evidence for the full 15.2M-parameter model. Their absolute MFD values use a simplified single-seed comparison pipeline and should be read only as within-table deltas, not as recomputations of the main or multi-seed results. A 7-configuration ablation (Appendix D and Appendix Figure 6) suggests sparsity loss is most important when zero-inflation exceeds ≈85%, KL divergence matters more on smaller datasets, and co-exclusion constraints are population-specific.

#### Ablation findings (593K model)

Removing sparsity loss causes the largest within-table MFD degradation on the compendium (+0.0292) but not AGP (+0.0000). Thus, this ablation supports the narrower claim that the explicit sparsity penalty is most useful under very high zero-inflation; the central AGP result should instead be attributed to the zero-inflated decoder, prevalence-aware initialization, and prevalence loss as a combined design. KL removal hurts AGP more than the larger compendium (+0.0065 vs. +0.0037), and removing AGP-derived co-exclusion constraints hurts AGP (+0.0045) but slightly helps the compendium (−0.0025), consistent with population-specific ecological networks [Sonnenburg and Sonnenburg, 2019]. The prevalence-aware bias initialization adapts to compendium sparsity (0.899 vs. 0.646 in AGP) without dataset-specific tuning.

### 5.6 Alpha Diversity Gap

A persistent alpha diversity gap (AGP: 3.15 vs. 2.88 real; compendium: 2.01 vs. 1.95) remains unresolved. Inspection of generated abundance profiles suggests that, conditional on presence, the log-normal decoder yields abundances that are too uniform across present taxa and under-represent the heavy-tailed dominance patterns seen in real samples. Temperature scaling and dominance regularization failed; heavier-tailed abundance parameterizations, normalizing-flow decoders, or hierarchical discrete latents are the most promising directions forward.

## 6 Limitations and Conclusion

### Limitations

SparseDOSSA2 achieves lower sparsity deviation than our method (0.7% vs. 1.4% in the main comparison), and MIDASim also passes the operational sparsity threshold. Our claim is therefore limited to deep generative modeling at parametric-level sparsity preservation, not absolute sparsity superiority. PERMANOVA remains able to distinguish generated from real AGP samples (*F* = 64.29), showing that generated communities are not distributionally indistinguishable from real ones. MFD and FMFD are susceptible to dense mean-averaging or feature-metric artifacts, so the full metric suite is necessary. The evaluation does not include zero-inflated VAE/GAN baselines, so the evidence isolates the value of explicit zero-inflation more strongly than the value of diffusion per se. The hyperbolic embedding component remains an unvalidated design choice, and the model ignores functional and temporal information, leaves a persistent alpha-diversity gap, and relies on co-exclusion constraints that appear population-specific.

We present, to our knowledge, the first deep generative framework to match parametric-level microbiome sparsity preservation through a zero-inflated decoder and prevalence-aware initialization: on AGP, the full model reaches 1.4% sparsity deviation in the main comparison and 2.6% ±0.5% across three random seeds. SparseDOSSA2 is stronger on sparsity deviation and Jaccard distance; MIDASim is stronger on ecological distance metrics overall. Our model achieves the best prevalence correlation (0.996) and competitive Bray–Curtis and UniFrac discrepancies. The narrow takeaway is that realistic deep microbiome synthesis requires absence structure to be built into the model rather than left as an emergent property.

## A Dataset Details

The American Gut Project and Human Microbiome Compendium are used as publicly available research resources. Users of the data and any released code should follow the access terms, citation requirements, and licenses specified by the original data providers.

## B Hyperparameters

## C Auxiliary Loss Definitions

For minibatch generated compositions 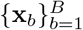, real minibatch compositions 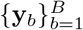, and latent encoder parameters (***µ***_*z*_, ***σ***_*z*_) when the comparison implementation uses a variational state, the auxiliary losses are:

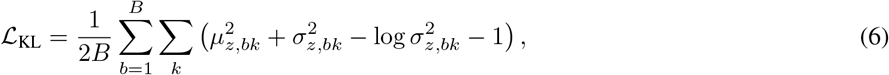

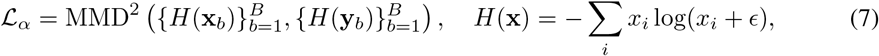

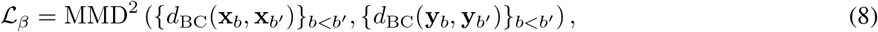

where *d*_BC_(**u, v**) = Σ_*i*_ |*u*_*i*_ *− v*_*i*_|*/* Σ_*i*_(*u*_*i*_ + *v*_*i*_) is Bray–Curtis dissimilarity and MMD uses an RBF kernel with the median heuristic. In configurations without a variational latent state, *ℒ*_KL_ is omitted.

## D Ablation Study Full Tables

## E Multi-Seed AGP Results

AGP experiments were run with 3 random seeds (42, 7, 123). Mean *±* std sparsity deviation is 2.6% *±* 0.5%. The per-seed deviations are: seed 42, 2.8%; seed 7, 1.9%; seed 123, 3.2%. These multi-seed AGP results support the central sparsity-preservation claim in the abstract and introduction. The ablation MFD values in Appendix D were not recomputed under this multi-seed protocol and therefore support only qualitative within-table sensitivity comparisons.

## F Computational Efficiency

All experiments run on a single NVIDIA A100 (40GB). Total compute for the project (including hyperparameter search and failed preliminary experiments) is approximately 200 GPU-hours.

## G Additional Biological Validation

**Figure 5:**
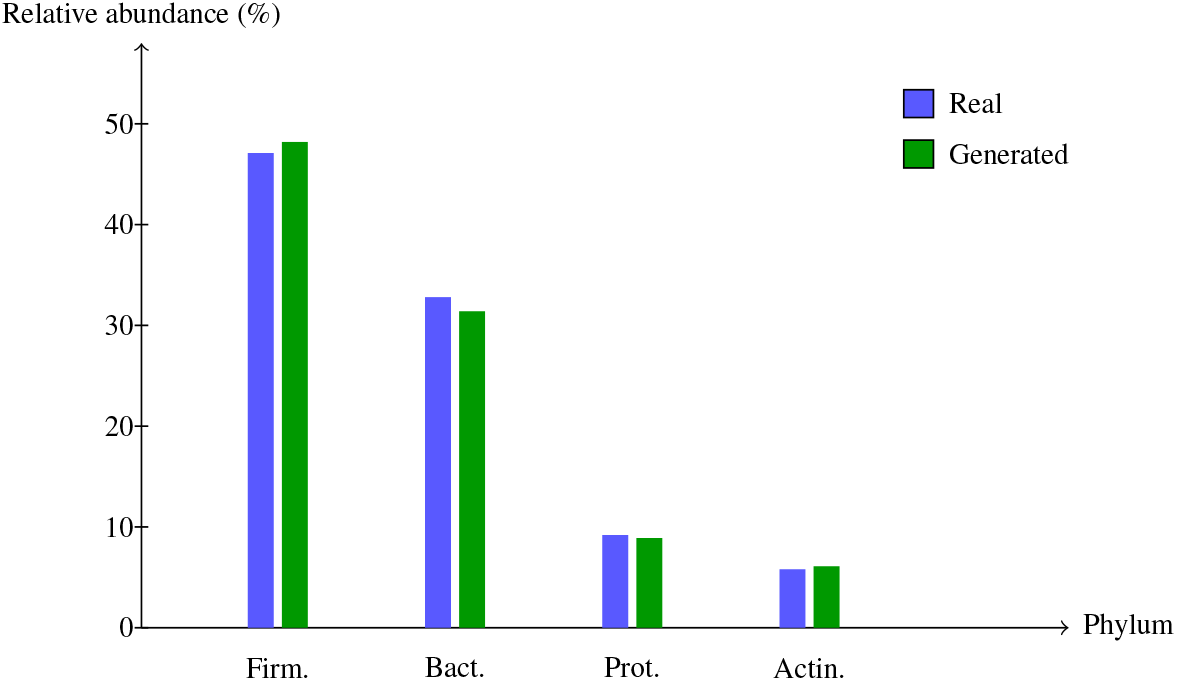
Biological validation as a TikZ grouped bar chart. Major phylum-level abundances are close between real and generated AGP samples: Firmicutes 47.1% vs. 48.2%, Bacteroidetes 32.8% vs. 31.4%, Proteobacteria 9.2% vs. 8.9%, and Actinobacteria 5.8% vs. 6.1%.

**Figure 6:**
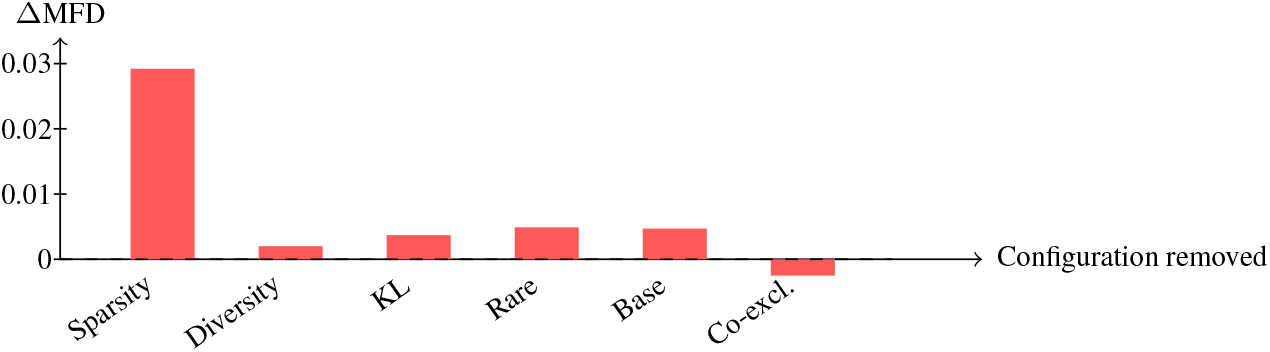
Ablation summary as TikZ bars for the non-Western compendium comparison model. Bars show within-table ΔMFD relative to the full 593K-parameter comparison model and should be interpreted only as diagnostic sensitivity results.

### Phylum-level composition

Firmicutes 48.2% (real: 47.1%), Bacteroidetes 31.4% (real: 32.8%), Proteobacteria 8.9% (real: 9.2%), Actinobacteria 6.1% (real: 5.8%).

### Pathogen detection

Clinically important pathogens (*Clostridoides difficile, Helicobacter pylori*) are correctly absent in generated samples from healthy donors, preserving clinical relevance.

### Diversity preservation

Beta diversity (Bray-Curtis) distribution matches real data with MMD = 0.015.

